# Intracellular biomacromolecule delivery by stimuli responsive protein vesicles loaded by hydrophobic ion pairing

**DOI:** 10.1101/2024.02.27.582187

**Authors:** Mikaela A. Gray, Alejandro de Janon, Michelle Seeler, William T. Heller, Nicki Panoskaltsis, Athanasios Mantalaris, Julie A. Champion

## Abstract

Therapeutic biomacromolecules are highly specific, which results in controlled therapeutic effect and less toxicity than small molecules. However, proteins and nucleic acids are large and have significant surface hydrophilicity and charge, thus cannot diffuse into cells. These chemical features render them poorly encapsulated by nanoparticles. Protein vesicles are self-assembling nanoparticles made by warming elastin-like polypeptide (ELP) fused to an arginine-rich leucine zipper and a globular protein fused to a glutamate-rich leucine zipper. To impart stimuli-responsive disassembly and small size, ELP was modified to include histidine and tyrosine residues. Additionally, hydrophobic ion pairing (HIP) was used to load and release protein and siRNA cargos requiring endosomal escape. HIP vesicles enabled delivery of cytochrome c, a cytosolically active protein, and significant reduction in viability in traditional two-dimensional (2D) human cancer cell line culture and a biomimetic three-dimensional (3D) organoid model of acute myeloid leukemia. They also delivered siRNA to knockdown protein expression in a murine fibroblast cell line. By examining uptake of positive and negatively charged fluorescent protein cargos loaded by HIP, this work revealed the necessity of HIP for cargo release and how HIP influences protein vesicle self-assembly using microscopy, small angle x-ray scattering, and nanoparticle tracking analysis. HIP protein vesicles have the potential to broaden the use of intracellular proteins for various diseases and extend protein vesicles to deliver other biomacromolecules.

## 1. Introduction

Intracellular protein and nucleic acid therapeutics could have high potency and specificity against diseases such as cancer by directly engaging in cell signaling pathways or metabolism.^1–3^ With these advantages, biomacromolecules have the potential to exhibit less toxicity to healthy tissue than small molecule drugs.^2,4,5^ However, there are significant delivery challenges for intracellular protein and nucleic acid therapeutics: instability following administration due to degradation by proteases/nucleases or unfolding, immunogenicity, and inability to cross cell membranes.^6,7^ Nanoparticles such as liposomes, polymeric nanoparticles, and protein nanoparticles encapsulate proteins and nucleic acids to protect them from degradation, can be decorated to allow for targeted delivery, improve endocytic delivery, and provide controlled release.^8–11^ Liposome and polymeric nanoparticles often use organic solvents during fabrication and some formulations possess toxicity related concerns. Organic solvents can denature or change the native structure of a protein by disrupting hydrophobic interactions, which leads to loss of protein function.^12^ One common challenge many nanoparticles face is poor encapsulation of large, charged biomacromolecule cargoes because most nanoparticles were developed for smaller and more hydrophobic cargo and their building blocks tend to be smaller than proteins. When encapsulating protein and nucleic acid cargos, many nanocarriers have low encapsulation efficiency (EE), undesired cargo leakiness, and an initial cargo burst release.^13,14^

To overcome the challenges associated with encapsulating biomacromolecule cargos, hydrophobic ion pairing (HIP) has been used, which reduces the solubility of compounds by forming reversible ionic interactions between a charged therapeutic and an oppositely charged hydrophobic counterion.^15^ HIP decreases electrostatic repulsion of the therapeutic by neutralizing the molecule’s natural surface charge and increases the hydrophobicity by coating the molecule’s surface with hydrophobic domains.^16^ HIP is reversed when the therapeutic and counterion are dissociated by competition from other ions due to change in pH or salt concentration. Nanoparticles are exposed to increasingly acidic environments from weakly acidic endosomes (pH 5.5-6.0) to more acidic lysosomes (pH 4.5-5.0) that cargo must usually escape to achieve cytosolic delivery.^17^ Sun and coworkers showed that pairing insulin and hydrophobic ion oleic acid in pH sensitive nanoparticles increased the encapsulation efficiency of insulin by more than 200%.^18^ HIP has been utilized to load and deliver small molecules, peptides, and proteins, including insulin, polymyxin B, and lysozyme, in various types of nanoparticles.^15^ HIP complexes improve membrane permeability because of their high solubility in the cell membrane and improve the oral bioavailability of many drugs.^19^ A similar strategy of increasing hydrophilic cargo hydrophobicity has been used in noncationic micelles to co-deliver a hydrophilic nucleic acid with a hydrophobic small molecule drug.^11^

We propose that self-assembling protein vesicles, which are genetically tunable, biodegradable nanoparticles made without solvent, could be used for cytosolic biomacromolecule delivery.^20^ Protein vesicles consist of a globular protein (mCherry in this work) attached to a glutamic acid-rich leucine zipper (Z_E_) that binds strongly to an arginine-rich leucine zipper (Z_R_) attached to an elastin-like polypeptide (ELP) to form a protein complex. Above the transition temperature (T_t_), ELP undergoes a conformational change from soluble to insoluble depending on salt concentration and type, Z_E_/Z_R_ molar ratio (always < 1), ELP sequence, and protein concentration.^21–26^ Our group has demonstrated doxorubicin HCl loading and release *in vitro* from covalently crosslinked protein vesicles.^27^ However, the crosslinking used to stabilize the vesicles in physiological conditions limits the size of cargo that can be released from protein vesicles. Recently, we rationally designed an ELP variant to produce stable protein vesicles at physiological salt concentration without crosslinking.^21^ This was accomplished by substituting tyrosines (Y) for valines (V) at guest positions (X) in the ELP sequence (VPGXG)_25_ (P = proline, G = glycine). Additionally, we have substituted histidines (H) for valines to impart pH-responsiveness, evidenced by vesicle disassembly in acidic conditions, to aid in escaping the endosome.^28^Taking these results together, we hypothesize that combining the pH-sensitive ELP and hydrophobic ELP will form new protein vesicles that stably assemble in physiological salt concentration with pH triggered disassembly (Table S1).

Encouraged by the breadth of therapeutics encapsulated into polymeric and lipid based nanoparticles using HIP in the literature and the ability of protein vesicles to encapsulate and deliver hydrophobic small molecules,^15,27,29^ this work examined how pH-sensitive protein vesicles can be modified and combined with HIP for encapsulation, release, and cytosolic delivery of charged protein and siRNA cargos. Two commonly used, generally regarded as safe (GRAS), anionic counterions were selected, oleic acid (OA) and sodium docusate (SD) (Figure S2A,B). Oleic acid is a fatty acid used in cholesterol supplements and hair products. It is also a known transdermal permeation enhancer.^30^ SD is a laxative that decreases surface tension to allow for water penetration into stool.^31^ In an oral delivery study of leuprorelin, insulin, and desmopressin peptides using HIP with self-emulsifying drug delivering systems, sodium docusate showed the best EE.^32^ Benethamine (BA) was selected as the cationic counterion (Figure S2C). Most cations are toxic, but BA is a lipophilic amine used as a drug salt in FDA approved penicillin G, a long acting penicillin. BA has been paired with retinoic acid for improved EE in a nanostructured lipid carrier compared to two other cationic counterions.^33^ Here, we develop a formulation of protein vesicles using engineered proteins and HIP to intracellularly deliver functional protein and siRNA cargo, expanding the cargo types used for protein vesicle delivery.

## 2. Results and Discussion

### 2.1. Protein vesicles made from rationally designed ELPs

For this work, a pH-sensitive ELP with 15 histidine residues (H_15_-Z_R_-ELP) and an ELP with 5 tyrosine residues (Y_5_-Z_R_-ELP) that makes vesicles stable in physiological salt were selected expressed as fusions with Z_R_ in *Escherichia coli* (*E. Coli*) and purified (Table S1, Figure S1). To visualize vesicles, model fluorescent protein mCherry-Z_E_ was selected as the globular protein, expressed as a fusion to Z_E_ in *E. Coli* and purified. The proteins were mixed in phosphate buffered saline (PBS) and warmed to 25 ºC for one hour to form vesicles (Figure 1A). A 12 Y_5_-Z_R_-ELP: 1 H_15_-Z_R_-ELP molar ratio was selected over 1:1 and 2:1 ratios as it formed vesicles with the smallest size and a measurable amount of H_15_-Z_R_-ELP without aggregation using a 0.05 Z_E_/Z_R_ molar ratio (Figure S3). As in previous work,^21^ increasing the Z_E_/Z_R_ molar ratio to 0.3 further reduced vesicle diameter to 141.2 ± 3.64 nm by dynamic light scattering (DLS) with a polydispersity index (PDI) of 0.291 ± 0.03 (Figure 1B). As the vesicles were too small to observe morphology by fluorescence microscopy, we examined the turbidity profile (Figure S4).The turbidity remained high after the initial increase as is characteristic of stable vesicles; coacervates, which are an intermediate phase in the vesicle assembly process display decreasing turbidity after the initial increase.^20,34^ Vesicle morphology was examined using a lower Z_E_/Z_R_ ratio (0.05), to make large structures visible using epifluorescence, where the particles were hollow and spherical (Figure 1C). The zeta potential of this formulation of protein vesicles was -9.76 ± 1.9 mV, approximately neutral to slightly negative. The vesicles maintained a stable diameter when diluted in PBS by 30% (the maximum for high enough concentration for DLS), showing that reduction in protein concentration did not induce disassembly (Figure 1B).

**Figure 1.**
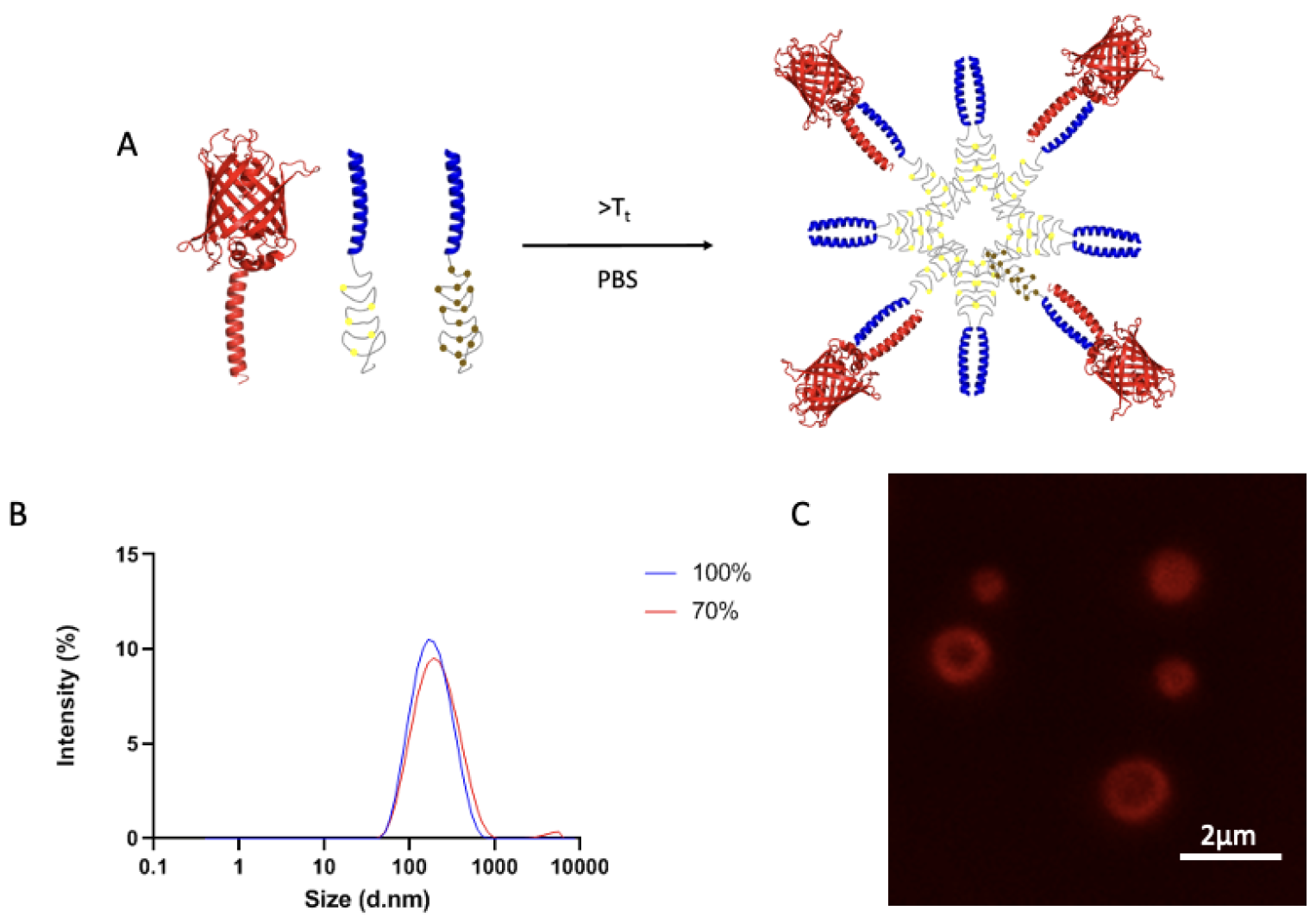
Protein vesicle characterization. ((A) Illustration of self-assembling protein building blocks composed of mCherry-Z_E_ and a mixture of Z_R_-ELPs using a 0.3 Z_E_/Z_R_ ratio, and 30 µM total ELP using a 12 Y_5_-Z_R_-ELP (labelled with yellow spheres):1 H_15_-Z_R_-ELP (labelled with brown spheres) molar ratio with 0.15 M NaCl. (B) DLS analysis showing size distribution of vesicles and stability of vesicles when diluted to 70% of original concentration into PBS. (C) Epifluorescence image of structures composed of mCherry-Z_E_ and a mixture of Z_R_-ELPs using a 0.05 Z_E_/Z_R_ ratio for enhanced visualization, and 30 µM total ELP using a 12 Y_5_-Z_R_-ELP: 1 H_15_-Z_R_-ELP molar ratio in 0.15 M NaCl. Image was digitally enlarged 10X.)

### 2.2. Protein vesicles deliver cytochrome c and siRNA loaded by HIP to cytosol

In order to determine if protein vesicles made from tyrosine and histidine ELPs can deliver a functional protein to the cytosol, cytochrome c (CytC) was selected as the cargo. Cytochrome C (CytC) induces apoptosis when localized in the cytosol of the cell.^35,36 37^ The charge ratios were calculated by using the molar ratio of protein and counterion and a surface potential calculator to determine the total protein surface potential relative to the counterion, which has 1 charge group.^38^ To determine if the amount of counterion influenced the size of protein vesicles, the concentration of CytC was kept constant while the amount of OA counterion varied (Figure S5A). Concentration of OA and protein vesicle size are linearly correlated, forming nano to micron sized particles as OA increased. The change in vesicle size could be due to hydrophobic interactions between the ELP and counterions influencing coacervate size.

Properties of HIP CytC OA or SD loaded vesicles are listed in Table 1 describing the encapsulation efficiencies, vesicle size, and PDI. Charge ratios of HIP OA or SD complexes listed in Table 1 resulted in no significant reduction in CytC fluorescence, demonstrating retention of protein structure (Figure S6A). After using HIP to load CytC into protein vesicles, vesicles were incubated with cells for 48 hours and cell viability was measured. First, we tested two charge ratios to determine for the effect on intracellular delivery. A charge ratio of 1 CytC: 1.8 OA did not result in intracellular delivery. However, a 1 CytC: 16 OA charge ratio yielded significant intracellular delivery in K562 acute myeloid leukemia cells (Figure S5B). Other researchers using HIP to load drugs into nanoparticles use significantly lower charge ratios, typically 1 cargo (small hydrophobic molecules, peptides, and proteins) :1 counterion, as higher charge ratios formed micelles in their complexes.^32,39–42^ We also observed less than 5% viability for 1 CytC: 16 OA or 1 CytC: 13 SD HIP loaded protein vesicles with HeLa cells (Figure 2C). OA has a higher LogP value (6.78) indicating greater hydrophobicity than SD (5.2), however both counterions enabled cytosolic delivery for charge ratios of 1 CytC:13 SD or 1 CytC:16 OA with these monovalent counterions. Vesicles loaded with CytC without HIP do not influence viability at all despite having a similar EE to HIP loaded vesicles. This demonstrates that HIP is changing the nature of cargo loading in vesicles, the ability of cargo to be internalized by cells, or the release of cargo from vesicles into the cytosol. Empty protein vesicles and HIP vesicles with non-toxic cargo (data in Section 2.2) do not reduce cell viability, confirming that the loss in viability is due to delivery of CytC not toxicity from delivery of hydrophobic ions.

**Table 1.** ((Molar and charge ratios of cargo to hydrophobic counterion, vesicle size, polydispersity index (PDI), and encapsulation efficiency (EE). n/m means values were not measurable. * indicates unpaired t-test comparison between no counter ion and HIP groups (* p< 0.05, ** p<0.01, *** p<0.001, or **** p<0.0001)))

| Cargo | Counterion | Molar Ratio | Charge Ratio | Vesicle Size [nm] | PDI | EE [%] |
| --- | --- | --- | --- | --- | --- | --- |
| CytC | none | - | - | 203.8 ± 8.08 | 0.315 ± 0.03 | 46.85 ± 5.82 |
|  | OA | 1: 127 | 1: 16 | 1155 ± 158*** | 0.123 ± 0.08* | 41.21 ± 2.46 |
|  | SD | 1: 108 | 1: 13 | 3485 ± 1130** | 0.194 ± 0.07* | 52.79 ± 3.90 |
| siRNA | none | - | - | 201.2 ± 2.73 | 0.552 ± 0.01 | 88.0 ± 3.72 |
|  | BA | 1: 3.7 × 10<sup>-10</sup> | 1: 1.8 × 10<sup>-11</sup> | 169.1 ± 1.06**** | 0.308 ± 0.01**** | 90.4 ± 3.00 |
| sfGFP(+10) | none | - | - | 261.8 ± 4.71 | 0.219 ± 0.01 | 64.09 ± 4.49 |
|  | OA | 1: 396 | 1: 35 | 174.1 ± 1.45*** | 0.263 ± 0.01** | 98.16 ± 1.43*** |
|  | SD | 1: 62 | 1: 5.4 | n/m | n/m | n/m |
| sfGFP(-10) | none | - | - | 216.96 ± 17.3 | 0.287 ± 0.05 | 16.52 ± 1.65 |
|  | BA | 1: 132 | 1: 14 | 193.9 ± 14.2 | 0.351 ± 0.05 | 20.37 ± 2.04 |

**Figure 2.**
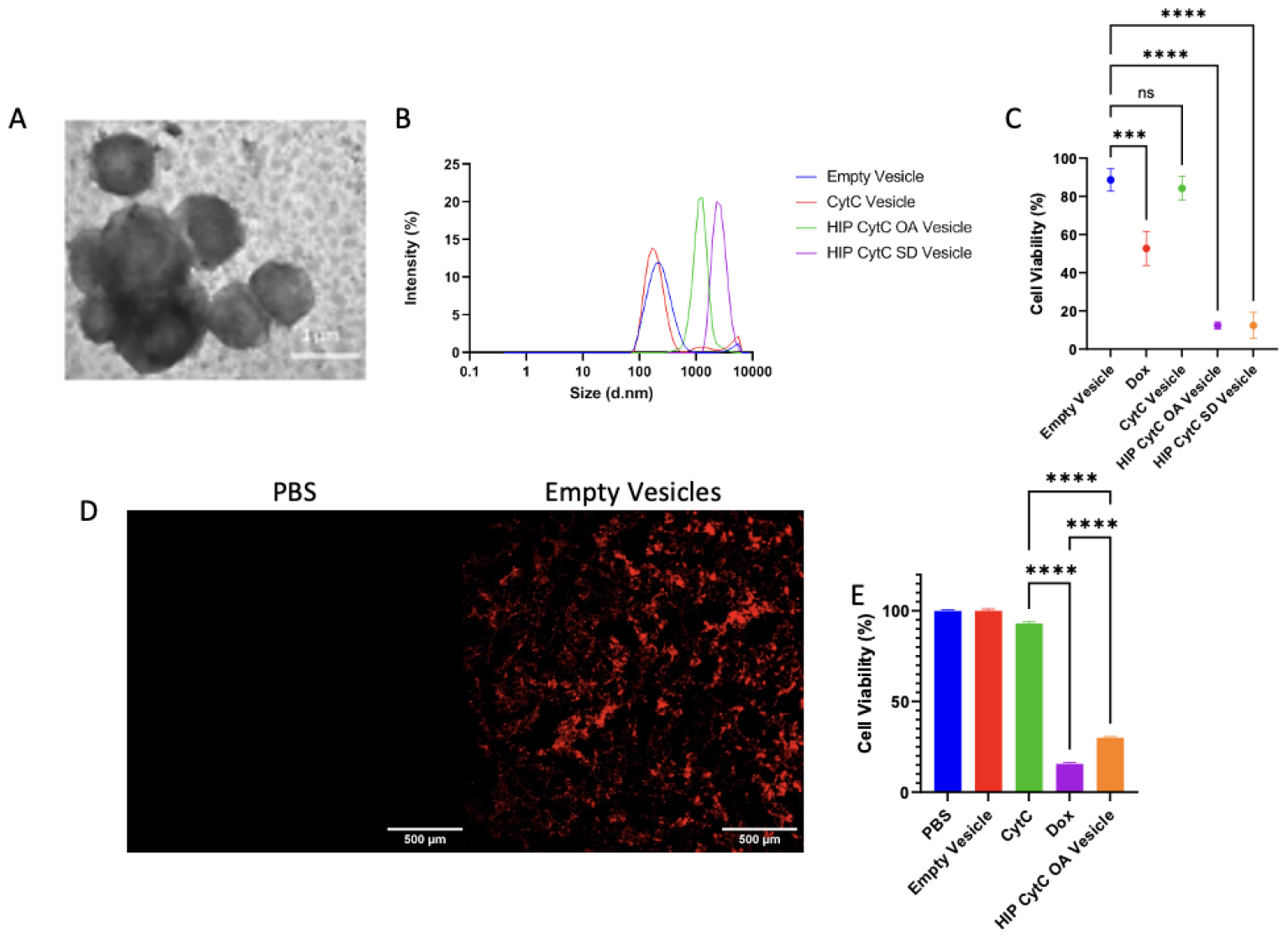
CytC vesicle characterization and delivery. ((A)TEM of HIP CytC OA loaded vesicles composed of mCherry-Z_E_ and a mixture of Z_R_-ELPs using a 0.3 Z_E_/Z_R_ ratio, 30 µM total ELP with a 12 Y_5_-Z_R_-ELP: 1 H_15_-Z_R_-ELP molar ratio, 10 µM CytC cargo, and 0.15 M NaCl. (B) DLS analysis showing size distribution of cargo-loaded vesicles. (C) HeLa cell viability measured by MTT assay 48 hours after treatment with 2 µM CytC relative to media control. (D) Vesicle diffusion within an acute myeloid leukemia organoid. (E) Acute myeloid leukemia bone marrow organoid K562 cell viability 24 hours after treatment with 2 µM CytC or 6 µM Dox showing ability to enter the cytosol of cells in a tumor-like environment. One-way ANOVA was utilized with p>0.05 n.s, p<0.001***, p<0.0001****, and n=3 groups with each experiment repeated at least twice. Error bars are standard deviation from the mean.))

To measure therapeutic delivery efficacy in a complex tumor-like environment, an acute myeloid leukemia organoid (using the K562 cell line) was utilized. Acute myeloid leukemia is a bone marrow cancer characterized by expansion and differentiation arrest of myeloid progenitor cells, with a 5-year survival of 24%.^43^ This biomimetic bone marrow organoid, containing collagen I, replicates the structural characteristics of human bone marrow and captures various biological cues unique to this tissue, exhibiting comparable porosity, pore size distribution, and stiffness, allowing cells to faithfully recreate spatial niches reminiscent of *in vivo* systems.^44^ Vesicles penetrated the 100-200 micron-sized niche pores within the organoid (Figure 2D).^44^ Vesicles delivered the cargo intracellularly resulting in less than 35% cell viability using a 2 µM dose of CytC after 24 hours of treatment (Figure 2E). This is significant because organoids have self-organization of acute myeloid leukemia niches and mirror leukemic metabolism yielding an environment more like human acute myeloid leukemia bone marrow tumor microenvironment compared to traditional 2D cell cultures.^44,45^

As delivery of a large hydrophilic protein using HIP vesicles resulted in intracellular delivery, we hypothesized that this technique would enable delivery of smaller hydrophilic nucleic acid cargo. After loading small interfering RNA (siRNA) designed to silence GFP, we characterized vesicle size, PDI, and encapsulation efficiency (Table 1). When a 1 siRNA: 3.7 × 10^−10^ BA charge ratio was utilized, the size of the vesicle and PDI reduced (Figure 3A). We tested intracellular delivery by quantifying GFP silencing using NIH3T3/GFP murine fibroblast cells 48 hours after treatment (Figure 3B, Figure S7). HIP siRNA BA vesicles resulted in greater than 70% GFP knockdown, like lipofectamine RNAiMAX (commercial gold standard), though 50% more siRNA cargo was used for vesicle delivery. However, HIP loaded vesicles delivered siRNA in serum supplemented media while lipofectamine RNAiMAX delivery required serum free media. Vesicles without HIP resulted in no GFP knockdown and we hypothesize that the siRNA interacted strongly with arginine’s in Z_R_ in the absence of HIP as cationic arginine residues have been used to condense anionic siRNA in other systems.^46,47^ Importantly, siRNA delivery by vesicles had no impact on cell viability while lipofectamine RNAiMAX reduced viability to less than 50% (Figure 3C). Altogether, this data demonstrates that HIP loaded mixed ELP vesicles are compatible for cytosolic delivery of two very different types of biomacromolecules.

**Figure 3.**
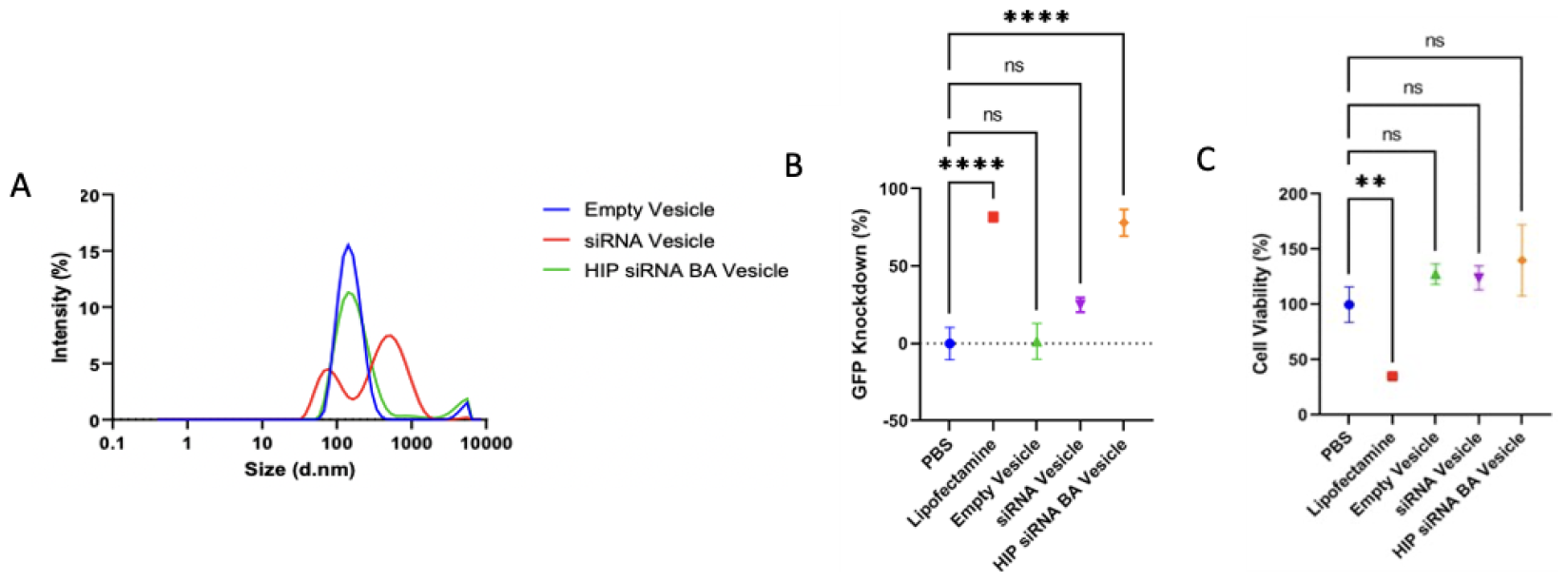
siRNA vesicle characterization and delivery. ((A) DLS analysis showing size distribution of cargo-loaded vesicles. (B) NIH3T3/GFP cell GFP knockdown relative to media control and lipofectamine positive control 48 hours after treatment. Lipofectamine RNAiMAX contained 100 nM siRNA and vesicle groups contained 150 nM siRNA. (C) NIH3T3/GFP cell viability measured by MTT assay 48 hours after treatment. One-way ANOVA was utilized with p>0.05 n.s, p<0.0001****, and n=3 groups with each experiment repeated at least twice. Error bars are standard deviation from the mean.))

### 2.2. Characterization of vesicle delivery of model positive and negative protein cargos

HIP has been used to increase polymer and lipid-based nanocarrier encapsulation efficiency by increasing cargo hydrophobicity.^15^ In order to characterize how HIP enables vesicle loading and cytosolic delivery of biomacromolecules, two model superfolder green fluorescent protein (sfGFP) cargos were utilized. Negatively and positively surface charged sfGFP model cargo proteins were expressed in *E. Coli* then purified (Figure S1).^34^ Cationic BA was used to pair with sfGFP(−10) and no loss of protein fluorescence, as a proxy for structure, was observed (Figure S6D). Vesicles loaded with sfGFP(−10) without HIP or with 1 sfGFP(−10):14 BA charge ratio HIP (to match CytC) had similar size, polydispersity index, and encapsulation efficiency (Table 1, Figure 4). Larger vesicles were made by decreasing the Z_E_/Z_R_ molar ratio to 0.05 in order to visualize the spatial distribution of the cargo in the vesicles, although there could be differences in smaller vesicles used for delivery due to tighter packing and smaller size with 0.3 molar ratio. HIP loaded sfGFP(−10) vesicles have cargo loaded within the lumen distributed away from the membrane, while sfGFP(−10) vesicles without HIP have cargo-membrane colocalization (Figure 4A). Additionally, the zeta potential of sfGFP(−10) vesicles was -8.93 ± 0.77 mV and sfGFP(−10) BA HIP vesicles was - 12.9 ± 1.0 mV. Despite the addition of cationic BA, sfGFP(−10) BA HIP vesicles have significantly more negative surface charge (p = 0.0055) than empty and sfGFP(−10) loaded vesicles. We hypothesize this could be due to less surface exposed positive Z_R_ from changes in vesicle packing and organization caused by hydrophobic HIP complex.

**Figure 4.**
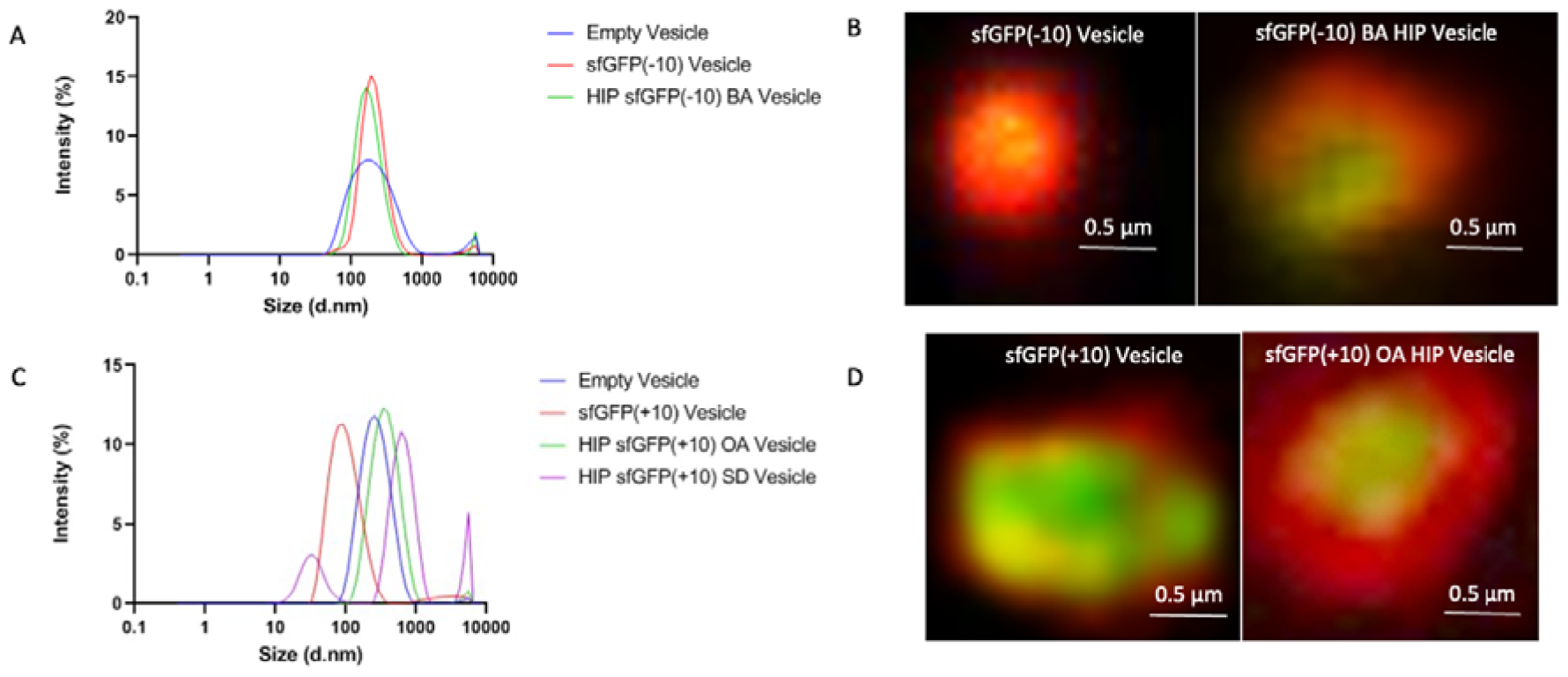
Empty, sfGFP(−10), and sfGFP(+10) loaded vesicle characterization ((A) DLS size distribution of cargo-loaded vesicles composed of mCherry-Z_E_ and a mixture of Z_R_-ELPs using a 0.3 Z_E_/Z_R_ ratio, 30 µM total ELP using a 12 Y_5_-Z_R_-ELP: 1 H_15_-Z_R_-ELP molar ratio with and without 10 µM of sfGFP(−10) cargo in 0.15 M NaCl or with HIP sfGFP(−10) BA. (B) Epifluorescence images of structures composed of larger vesicles using a 0.05 Z_E_/Z_R_ ratio (all other conditions the same as (A for cargo visualization). Scale bars are 0.5 µm and images were digitally magnified 100X. (C) DLS size distribution of cargo-loaded vesicles composed of mCherry-Z_E_ and a mixture of Z_R_-ELPs using a 0.3 Z_E_/Z_R_ ratio, and 30 µM total ELP using a 12 Y_5_-Z_R_-ELP: 1 H_15_-Z_R_-ELP molar ratio with and without 10 µM of sfGFP(+10) cargo in 0.15 M NaCl or with HIP sfGFP(+10) OA or SD. (D) Epifluorescence images of structures composed of larger vesicles using a 0.05 Z_E_/Z_R_ ratio (all other conditions the same as (A for cargo visualization). Scale bars are 0.5 µm and images were digitally magnified 100X.))

The same vesicle fabrication and characterization was performed for sfGFP(+10) loaded vesicles using OA and SD. Even with a lower charge ratio (1 sfGFP(+10): 5.4 SD) HIP sfGFP(+10) loading with SD resulted in unstable aggregates (Table 1), though sfGFP(+10) fluorescence was not impacted (Figure S6C). Protein vesicles loaded with sfGFP(+10) without HIP and 1 sfGFP(+10): 35 OA charge ratio HIP were stable with diameters of 261.8 ± 4.71 nm and 174.1 ± 1.45 nm, respectively (Table 1). A higher charge ratio of OA was used for sfGFP(+) than CytC to maintain vesicle sizes in the same range as both sfGFP(+10) without HIP and sfGFP(−10) with BA HIP for the delivery comparison experiments. HIP loading of sfGFP(+10) with OA did result in a loss of less than 30% of sfGFP(+10) fluorescence (Figure S6). Using this ratio of OA to load sfGFP(+10) into vesicles significantly improved the encapsulation efficiency by transiently increasing the cargo hydrophobicity (Table 1). The location of the sfGFP(+10) cargo changed, as with the sfGFP(−10) vesicle, when using HIP (Figure 4D).

To perform more detailed structural characterization of HIP vesicles, sfGFP(−10) and sfGFP(−10) BA HIP vesicles were selected because of cargo fluorescence tracking ability and BA is more easily dissolved than the anionic counterions. We first characterized the effect of cargo loading on ELP properties. ELPs are thermo-responsive and exhibit lower critical solution temperature phase behavior when heated. This behavior is characterized by the transition temperature, T_t_, defined as the inflection point of the turbidity profile as a function of temperature. By measuring the T_t_ of empty, sfGFP(−10) loaded, and sfGFP(−10) BA HIP loaded vesicles, we found vesicles undergo up to three transitions (Figure 5). Two are seen for all vesicles, related to the two different ELPs used in this work,^21,28^ and the third is only observed for HIP loaded vesicles. When multiple ELPs are present in a solution, there are multiple transition temperatures.^48^ Empty and sfGFP(−10) vesicles have the same T_t_ values, indicating the sfGFP(−10) cargo does no impact T_t_. Conversely, the first T_t_ of sfGFP(−10) BA HIP vesicles is lower T_t_. BA is a hydrophobic salt that is likely responsible for the depression as both hydrophobicity and salt concentration influence the T_t_.^21,34^ HIP-cargo complexes used in this work include counterions above their respective critical micelle concentrations for pure counterion solutions. This could explain the observed third T_t_, although we do not detect micelles by DLS or SAXS.

**Figure 5.**
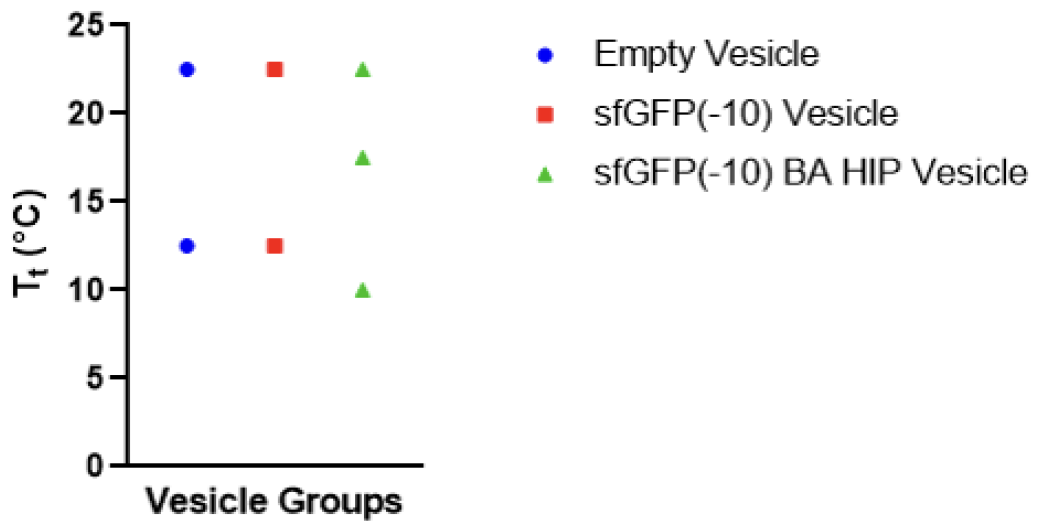
Effect of cargo and HIP on vesicle transition temperatures, T_t_. (Transition temperatures of empty, sfGFP(−10), and sfGFP(−10) BA HIP vesicles with 10 µM sfGFP(−10), were determined from the midpoint of each linear region of turbidity profile.)

To further analyze changes in vesicles with and without cargo and HIP, small angle x-ray scattering (SAXS) was employed to characterize empty, sfGFP(−10) loaded, and sfGFP(−10) BA HIP loaded vesicles formed in PBS at 25 ºC then adjusted to pH 7.4 or pH 6 and incubated for 2 or 4 hours before analyzing structures for 2 hours per sample. Empty vesicles were best modeled using a combination of vesicle and cylinder models (Figure S9). The membrane thickness increased, and the cylinder (the shape of the protein amphiphiles) radius decreased at pH 6, suggesting that the ELP was relaxing or extending as histidine protonation disrupted hydrophobic interactions (Table 2). This agrees with our previous work using vesicles made from H_15_-Z_R_-ELP, where SAX analysis revealed that the radius of gyration (R_g_) of soluble protein increased at pH 5.5, indicating more extended, hydrophilic soluble protein released from vesicle disassembly.^28^ However, vesicles loaded with sfGFP(−10) or sfGFP(−10) BA HIP scatter as fundamentally different objects than empty vesicles due to the contents in the lumen. They were fit with core-shell-sphere and monodisperse Debye gaussian coil (MGC) models, indicating that the cargo changes the core scattering length density (Figure S9). sfGFP(−10) loaded vesicles demonstrated decreased membrane thickness at pH 6 and sfGFP(−10) BA HIP vesicles exhibited no change in membrane thickness. Vesicles loaded with sfGFP(−10) have a significant increase in scaling for the MGC model, which represents soluble protein, at pH 6 compared to pH 7.4 (Table 2). This suggests release of cargo or vesicle proteins due to the start of vesicle disassembly. Conversely, sfGFP(−10) BA HIP loaded vesicles exhibited a trend of increased core-shell-sphere scaling at pH 6 at 4-6 hours.

**Table 2.**
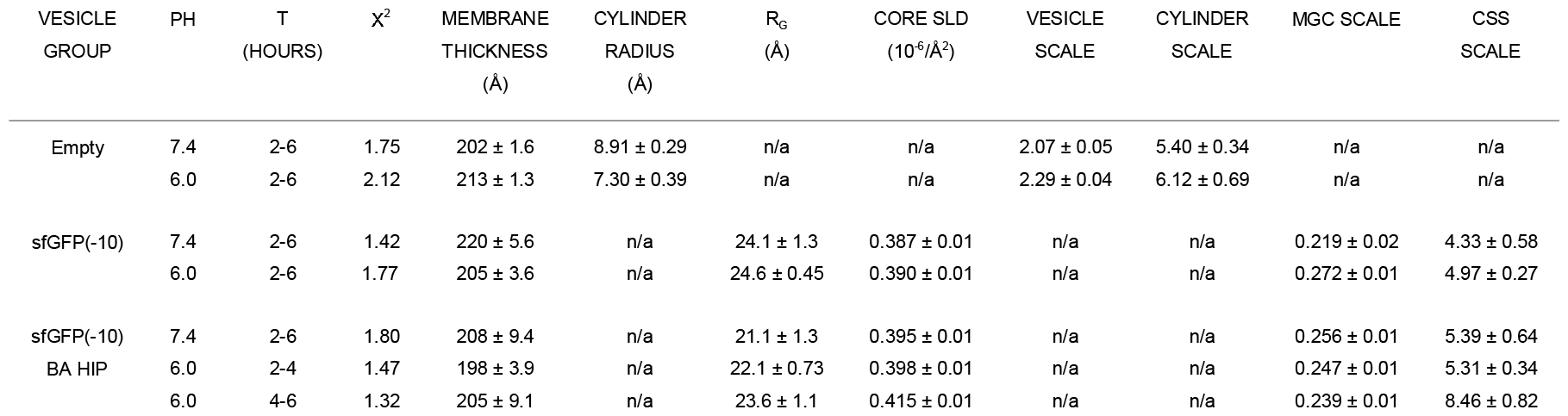
((SAXS model fitting parameters for empty, sfGFP(−10) loaded, and sfGFP(−10) BA HIP loaded vesicles. Empty vesicles were fit using vesicle and cylinder models, sfGFP(−10) and sfGFP(−10) BA HIP vesicles were fit using core shell and monodisperse Debye gaussian coil (MGC) models. T represents time in hours, X^2^ represents chi squared - a measure of model fitting, R_G_ represents the radius of gyration, Core SLD represents the core scattering length density, Vesicle Scale represents the scaling of that model, Cylinder Scale represents the scaling of that model, MGC Scale represents the scaling of that model, and CSS Scale represents the scaling of the core-shell-sphere model.))

Other samples (empty or sfGFP(−10) vesicles at all pHs, or sfGFP(−10) HIP vesicles at pH 7.4) exhibited no change between 2-6 hours. Nanoparticle tracking analysis showed that at pH 6, sfGFP(−10) HIP loaded vesicles and sfGFP(−10) loaded vesicles both had an order of magnitude less particles indicating vesicle disassembly (Figure S10). Ristroph and coworkers loaded polymixin B oleate HIP complexes into polymeric nanoparticles in water and found that at higher charge ratios (1:4), the nanoparticles have lamellar structures within the core that rearranged into an inverse hexagonal phase upon exposure to ions in PBS.^49^ At lower charge ratios (1:1) no change in assembly occurred. We hypothesize that a similar effect could occur and vesicle structure changes in part due to a change in HIP structure upon change in pH given the high charge ratios used to load vesicles. We hypothesize that the increase in core-shell-sphere scaling could result from the release of HIP cargo from disassembling vesicles resulting in less vesicles, but more core-shell-spheres, which are below the nanoparticle tracking analysis size limit of 30 nm.

### 2.3. Delivery of sfGFP(−10) and sfGFP(+10) by vesicles

In order to measure the influence of HIP loaded cargos on viability, characterize uptake, and study membrane interactions we used sfGFP(+10) and sfGFP(−10) to test anionic and cationic counterions, respectively. Vesicles with model cargos were delivered to HeLa cells for 24 hours at a dose of 1 µM cargo. The mass of vesicles was adjusted to account for differences in EE was using these mass ratios: 1 sfGFP(+10) OA HIP: 0.653 sfGFP(+10): 0.207 sfGFP(−10) BA HIP: 0.168 sfGFP(−10). sfGFP(−10) vesicle and sfGFP(−10) BA HIP vesicle groups resulted in significant sfGFP(−10) uptake (Figure 6A). No protein vesicle loaded with sfGFP(−10) had any influence on HeLa cell viability demonstrating that BA complexed with sfGFP(−10) in vesicles results in delivery without toxicity (Figure 6B). Using HIP to encapsulate sfGFP(+10) improved the EE significantly (Table 1) and sfGFP(+10) OA HIP loaded vesicles resulted in significantly higher HeLa cell uptake, while soluble sfGFP(+10) or vesicles loaded with sfGFP(+10) without HIP did not deliver any cargo (Figure 6C). Compared to sfGFP(−10) HIP vesicles, sfGFP(+10) HIP vesicles had a greater improvement on intracellular delivery, which could be due to the 2.5x higher charge ratio. However, CytC was delivered effectively with the same charge ratio used for sfGFP(−10), so there could also be effects from differences in cargo charge and size. As with vesicles delivering sfGFP(−10), no protein vesicles carrying sfGFP(+10) had any influence on HeLa cell viability. This shows that OA complexed to sfGFP(+10) in vesicles improves delivery without toxicity (Figure 6D). It also serves as a control for CytC delivery experiments, demonstrating that CytC exerted the toxic effect in Figure 2, not OA.

**Figure 6.**
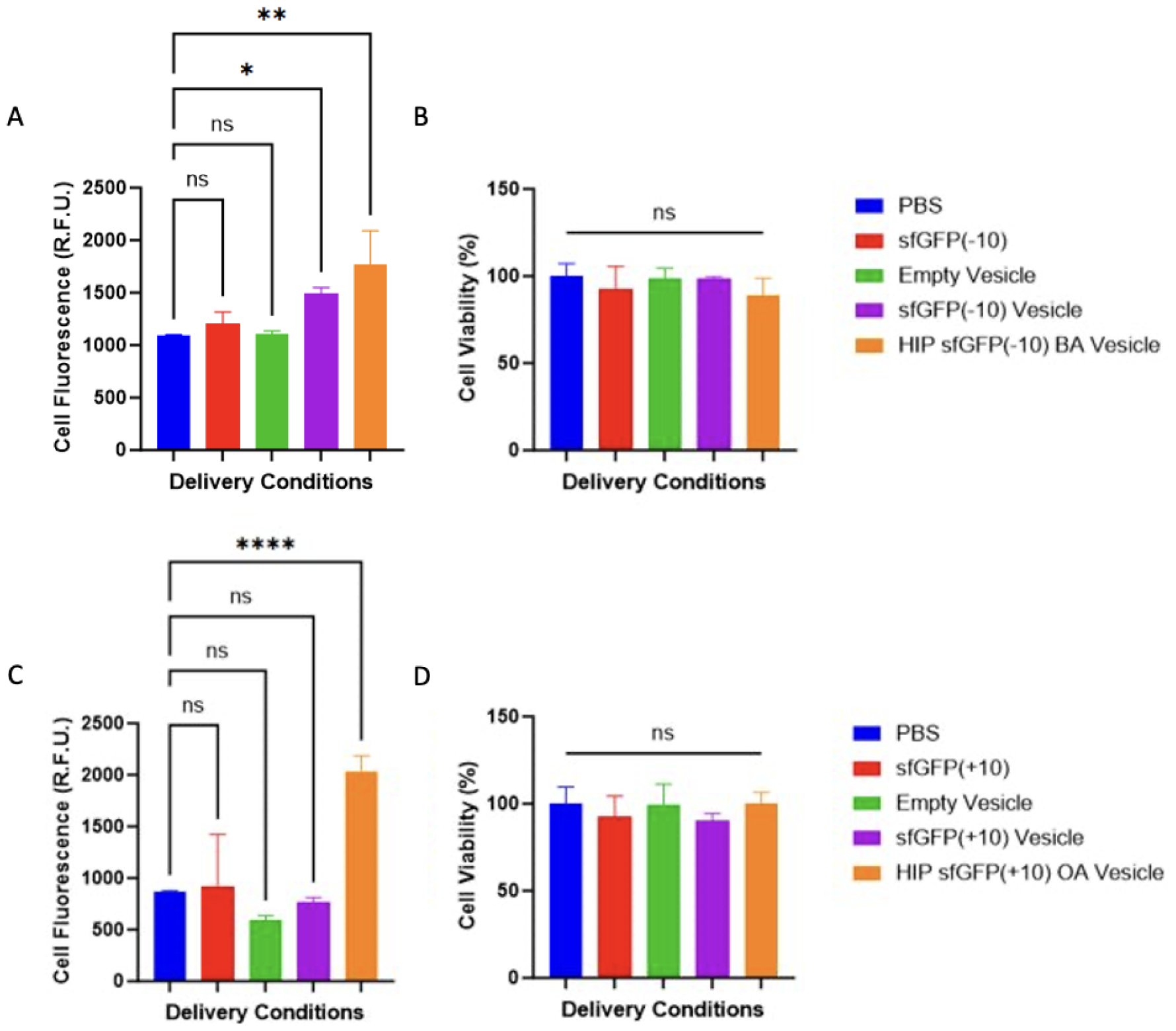
sfGFP(−10) and sfGFP(+10) loaded vesicle uptake and cellular viability ((A) Flow cytometry analysis showing median cell fluorescence of HeLa cells incubated with 1 µM sfGFP(−10) cargo fro 24 hours. (B) Viability 24 hours after treatment using MTT assay relative to media control. (C) Flow cytometry analysis showing median cell fluorescence of HeLa cells incubated with 1 µM sfGFP(+10) cargo fro 24 hours. (D) Viability 24 hours after treatment using MTT assay relative to media control. One-way ANOVA was utilized with p>0.05 n.s, p<0.05*, p<0.01**, p<0.0001****, and n=3 groups with each experiment repeated at least twice. Error bars are standard deviation from the mean.))

To determine how vesicles deliver cargo into the cytosol, we measured uptake at 37 and 4°C. Uptake was completely inhibited at 4°C, indicating an energy dependent uptake mechanism such as endocytosis (Figure 7A). We incubated vesicles with red blood cells (RBCs) to identify any vesicle membrane interactions, particularly at acidic pH found in endosomes. RBCs are not endocytic so any cargo or vesicle fluorescence increase in RBCs indicates membrane binding or disruption. When pH responsive vesicles disassemble at low pH, they fuse with another before completely disassembling.^28^ It is possible they could fuse with a membrane in a similar manner. Lipid nanoparticles merge with endosomal membranes in order to deliver cargos into the cytosol and exhibit improved cargo delivery when membrane fusion increases by use of ionizable lipids or cholesterol.^50,51^ Empty vesicles exhibited enhanced vesicle membrane-RBC interaction (mCherry fluorescence) at pH 5 compared to pH 7.4 (Figure 7B) suggesting that pH induced destabilization of vesicles increased membrane interactions. Conversely, neither of the cargo loaded vesicles had a pH dependent change in RBC interaction measured by mCherry fluorescence. It is possible that 1 hour incubation was not sufficient to destabilize cargo loaded vesicles and SAXS data indicated that empty vesicles respond differently to acidification that loaded vesicles. However, RBCs exposed to sfGFP(−10) loaded vesicles without HIP exhibited sfGFP fluorescence at pH 7.4 that dropped significantly at acidic pH. This could indicate that while the vesicle-RBC interactions were pH insensitive for cargo loaded vesicles, the interaction between the vesicle and cargo itself was pH sensitive without HIP and may have promoted release of sfGFP(−10) from vesicles.

**Figure 7.**
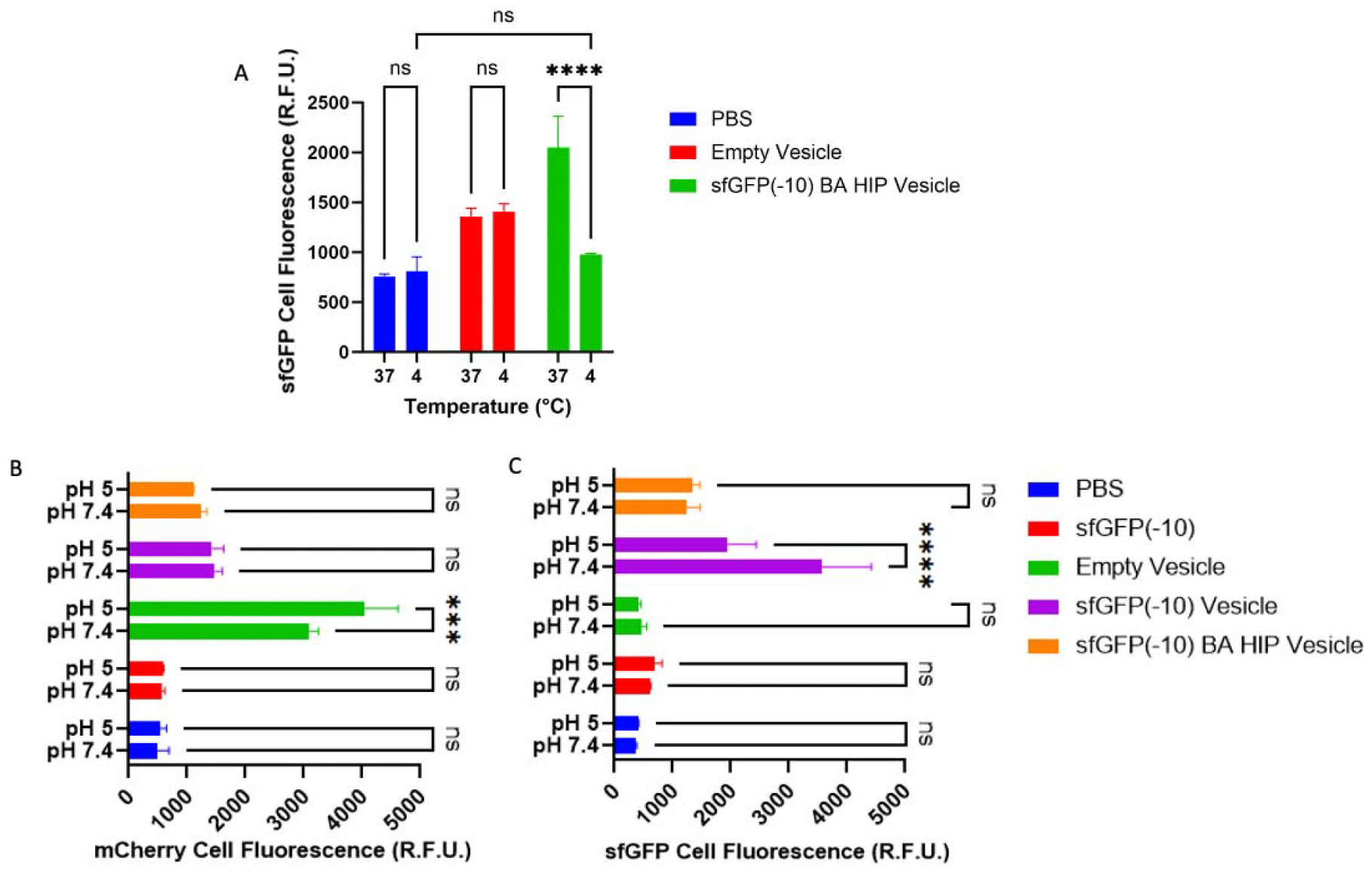
Energy dependent endocytosis and red blood cell membrane interactions of sfGFP(−10) loaded vesicles ((A) Energy dependent HeLa cell uptake using flow cytometry analysis of median fluorescence after 24 hours of incubation with 1 µM sfGFP(−10). (B, C) Red blood cell median fluorescence due to binding interactions with vesicles (B, mCherry fluorescence) or cargo (C, sfGFP fluorescence) at physiological and endosomal pH. 0.5% v/v RBC were treated for 1 hour, washed, and analyzed using flow cytometry. Vesicles were composed of mCherry-Z_E_ and a mixture of Z_R_-ELPs using a 0.3 Z_E_/Z_R_ ratio and 30 µM total ELP with 12 Y_5_-Z_R_-ELP: 1 H_15_-Z_R_-ELP molar ratio and 10 µM sfGFP(−10) cargo in 0.15 M NaCl, then treated according to the EE (using 0.413 or 0.51 mg/mL total vesicle proteins) to HeLa or RBC at 1 µM sfGFP. 2-way ANOVA was utilized with p>0.05 n.s, p<0.001***, p<0.0001****, and n=3 groups with each experiment repeated at least twice. Error bars are standard deviation from the mean.))

## 3. Conclusion

In this work, a mixed ELP protein vesicle formulation, where one ELP imparts increased stability and the other yields pH-responsiveness, was developed and characterized for biomacromolecule cargo delivery. HIP loaded vesicles enabled improved uptake of both positively and negatively charged proteins and siRNA without influencing cellular viability by using counterions with one charge group. We demonstrated that vesicles can deliver CytC into the cytosol when using HIP above a critical charge ratio, resulting in a significant reduction in cancer cell viability. Likewise, HIP loaded vesicles delivered siRNA in the presence of serum without decreasing cell viability. HIP loading of cargo was essential to successful delivery by protein vesicles and was observed to change the self-assembly of vesicles and the cargo location within the vesicle while also affecting cargo-membrane interactions at acidic pH. The combination of protein vesicles and HIP loading broadened the types of cargos delivered using protein vesicles and future work can explore the delivery of more cargo types both *in vitro* and *in vivo*.

## 4. Experimental Section/Methods

### Expression and Purification of Fusion Proteins

The pET28a-sfGFP-His plasmids with charged sfGFP variants were purchased from Genscript. sfGFP variants were expressed in *E. Coli* BL21(DE3) star. To express these proteins, 1 L of lysogeny broth (LB) containing 50 mg kanamycin was inoculated with 5 mL of overnight culture at 37 °C then induced with 1 mM isopropyl ß-D-1 thiogalactopyranoside (IPTG) when the optical density at 600 nm was greater than 0.7. After 5 hours of expression, cultures were collected by centrifugation at 4000 g for 10 minutes. The pellets were resuspended in lysis buffer containing 300 mM NaCl, 50 mM NaH_2_PO_4_, and 10 mM imidazole and lysed by sonication. Next, the supernatant was collected after centrifugation at 10,000 relative centrifugal force (rcf) for 10 minutes and incubated with Ni-nitrilotriacetic acid (NTA) agarose resin (Qiagen) for at least 1 hour at 4°C. This mixture was loaded in an Econo-Column (Biorad), washed with 100 mL of 25 mM imidazole, and 10 mL of elution was collected using 250 mM imidazole. Purity was verified using sodium dodecyl sulfate polyacrylamide gel electrophoresis (SDS-PAGE), then the proteins were buffer exchanged into phosphate buffered saline (PBS) by dialysis using a 10 kDa MWCO membrane with three buffer exchanges at 4 °C.

The genes for H_15_-Z_R_-ELP and Y_5_-Z_R_-ELP were purchased from Genscript. mCherry-Z_E_ was created previously.^52^ These three proteins were expressed in AF-IQ BL21 (DE3) *E. coli* in 1 L of LB containing 200 mg ampicillin and 34 mg chloramphenicol and collected using centrifugation at 4000 rcf for 10 minutes. mCherry-ZE was purified using the same method as sfGFP. Z_R_-ELPs were purified by affinity chromatography with the same resin and columns as above. Cells were lysed under denaturing conditions in 8 M urea, 100 mM NaH_2_PO_4_, and 10 mM TrisCl at pH 8, washed with 8 M urea at pH 6.3, and eluted into 6 M guanidine hydrochloride, 100 mM NaH_2_PO_4_, and 10 mM TrisCl, as in our past work.20,27,28,34 Y_5_-Z_R_-ELP purification used 5% n-octyl-ß-D-glucopyranoside in the lysis buffer as in our past work.21 After purification, the protein was buffer exchanged into miliQ water and freeze dried for long-term use.

### HIP Complex and Vesicle Formation

HIP complexes were formed by a simple mixing method. Cargo proteins at a concentration of 1 mg/ml or 50 µM siRNA were mixed with varying concentrations of counterion at equal volumes on ice before being added to vesicle proteins, the conditions used for each complex are in Table 1. Vesicle solutions were mixed on ice starting with miliQ water, 30 µM Z_R_-ELPs, mCherry-Z_E_, cargo, then 10X PBS to achieve a final salt concentration of 0.15 M. Unless otherwise stated, a Z_E_/Z_R_ ratio of 0.3 was utilized. The amount of cargo added was optimized based on the encapsulation efficiency so that the maximum loading was achieved (depending on the cargo concentration was between 6 and 10 µM).

### Characterization Techniques for Vesicles

Turbidity was determined by measuring absorbance at 400 nm with a Biotek Instrument Synergy H4 Hybrid multimode microplate reader set to 25 °C as the ice cold solution warmed over an hour. Size and zeta potential were measured by dynamic light scattering (DLS) using Malvern Instruments Zetasizer NanoZS with a 4 mW He-Ne laser at 633 nm wavelength to detect backscattering (173°) using PBS at pH 7.4 solvent conditions and protein material selection for DLS or 0.1X PBS at pH 7.4 solvent conditions and protein material selection for zeta potential. Z-average values were used for vesicle size. Electrophoretic mobility was converted to zeta potential using the Smoluchowski approximation. Encapsulation efficiency of protein vesicles was determined using 100 kDa MWCO 1 mL centrifugal filters to separate vesicles from unencapsulated cargo by loading vesicles in PBS then centrifuging at 12,500 rcf for 10 minutes. Fluorescence and absorbance calibration curves were made for each sfGFP charge variant and CytC using the same microplate reader used for turbidity measurements. Filtrate and retentate fluorescence or absorbance were measured to calculate EE.

Fluorescence imaging of vesicles was performed using an epifluorescence microscope (Zeiss Axio Observer Z1) using a 100X oil immersion lens. For high magnification imaging, transition electron microscopy (TEM), using a JEOL 100CX-II and imaged at 100 kV, grids were prepared using 10 µL sample on copper grids, letting the sample adhere for 5 minutes, washing in water for 30 seconds, staining with 1% phosphotungstic acid for 10 seconds, washing again for 30 seconds, and drying overnight. Small Angle X-Ray Scattering (SAXS) data was collected using a Rigaku BioSAXS 2000 instrument operated by the Center for Structural Molecular Biology at the Oak Ridge National Laboratory. The wavelength was 1.542 Å using momentum transfers 0.008 < Q < 0.70 Å^-1^. SAXSLab 4.0.2 software corrected for solvent background scattering, then data analysis was performed using SasView software (https://sasview.org). Data between 0.008 and 0.35 Å^-1^ was fit. Nanoparticle tracking analysis data was collected using a Nanosight NS300.

### Cell Culture and Cargo Uptake

HeLa cells (ATCC) were seeded at 15,000 cells/well using Dulbecco’s modified Eagle medium (DMEM) supplemented with 10% fetal bovine serum (FBS) in 96 well plates for 24 hours prior to treatment, washed with PBS, then treated with sfGFP or CytC loaded vesicles and controls in media containing DMEM and 10% FBS at 37°C in a humidified environment containing 5% CO_2_. K-562 (human erythromyeloblastoid leukemia cell line ATCC, UK; CCL-243), originally derived from a patient with blast crisis (acute leukemia phase) of chronic myeloid leukemia, was chosen to model acute myeloid leukemia. For 2D experiments, cells were seeded at 25,000 cells/well in 96 well plates for 24 hours prior to treatment in Roswell Park Memorial Institute 1640 Medium (RPMI; Corning, Manassas, VA) supplemented with 10% v/v fetal bovine serum (FBS, heat inactivated; Gibco, NY), washed with PBS, then treated with vesicles in supplemented media at 37°C in a humidified environment containing 5% CO_2_ for 48 hours. For 3D experiments, 100 µl of cell suspension (4 × 10^6^ cells/scaffold) were seeded onto the scaffolds, placed in 24-well tissue culture plates and incubated for 30 min at 37 ºC in an atmosphere of 5% CO_2_ to allow cell adhesion prior adding 1.5 ml of culture medium. Scaffolds were cultured for 7 days with full media exchange every other day prior adding the vesicles. NIH 3T3 fibroblast cells expressing eGFP (Cell BioLabs) were seeded at 35,000 cells/well in a 96 well plate for 24 hours before treatment using DMEM supplemented with 10% FBS. All groups besides lipofectamine RNAiMAX were treated in the presence of serum containing media and cultured at 37 ºC in an atmosphere of 5% CO_2_.

### Scaffold Preparation

Scaffolds were fabricated by thermally induced phase separation as previously described.^44,53,54^ Briefly, polyurethane pellets (Estane 58300; Velox, Germany) were dissolved in dioxan (99.8% pure, Merck Millipore, Germany) to form a polymer solution (5 wt%) that was frozen at -80 C with subsequent sublimation in an ethylene glycol bath at - 15 °C. The scaffolds (pore size 100-250 um, porosity 90-95%) were cut in 5 × 5 × 5 mm cubes and pre-wetted in ethanol (70% v/v, Thermo Scientific) for 1 minute, then washed in PBS for 20 min and centrifuged for 10 min at 2500 rpm in PBS. Scaffolds were coated with collagen type I from calf skin (Sigma-Aldrich). Cubes were transferred into a 62.5 µg/ml collagen solution solublized in 0.1 M acetic acid and dissolved in deionized water at pH 7 (re-adjusted with addition of 0.1 M NaOH) and centrifuged at 2000 rpm for 20 min in the protein solution followed by a final centrifugation step in PBS at 1500 rpm for 10 min to unblock the surface pores. Scaffolds were sterilized by UV sterilization (8 min exposure at 230 v, 50 Hz, 0.14 A, UV lamp, Thermo Fisher) and 2 hours immersion in ethanol (70%). The collagen-coated scaffolds were washed twice for 15 min in PBS before adding media and placing in a humidified incubator for 3 days at 37 °C and 5% CO_2_ prior to seeding with leukemia cells.

### Cargo Internalization

Following incubation with anti-GFP siRNA loaded vesicles or sfGFP loaded vesicles and controls, cells were detached using trypsin and resuspended in PBS with trypan blue to quench extracellular fluorescence. The green fluorescence of 10,000 cells was measured using 488 nm excitation on the CytoFLEX flow cytometer (BD Bioscience). Gating was done on PBS groups to isolate single cells and compare increase or decrease in GFP signal (Figure S11).

### Cell Viability

The *in vitro* toxicity of vesicles containing the different cargos was quantified using a MTT assay. After vesicle incubation, the media was removed, cells were washed with PBS, then fresh media with 10 µL of 5 mg/ml MTT solution (Biotium) was added. The cells were incubated for 4 hours at 37°C in a humidified environment containing 5% CO_2_. Then, 200 µL of dimethyl sulfoxide (DMSO) was added to dissolve the formazan crystals. Absorbance of solutions was measured at 570 nm and 630 nm. Cell viability was calculated by using an absorbance ratio in a treated group relative to the PBS control group. Similarly, an MTS assay was performed on scaffolds, which is analogous to an MTT assay minus the solubilization step.

### Endosomal Membrane Interactions

Washed 5% turkey red blood cells (RBCs) from Fisher Scientific were diluted to a 0.5% v/v concentration in PBS. 100 µL of diluted nanoparticles were incubated with 10 µL RBC at 37°C for 1 hour in a 96 well plate in a humidified environment containing 5% CO_2_. For membrane binding experiments, 100 µL of RBCs were plated then treated with nanoparticles for 1 hour at 37°C in a 96 well plate in a humidified environment containing 5% CO_2_. After incubation with treatment groups, the RBCs were washed with PBS then analyzed using flow cytometry to measure mCherry or sfGFP signal as described above.

## Supporting Information

Supporting Information is available from the Wiley Online Library or from the author.

## Acknowledgements

The authors acknowledge financial support from the National Science Foundation BMAT Award 2104734. This work was performed in part at the Georgia Tech Institute for Electronics and Nanotechnology, a member of the National Nanotechnology Coordinated Infrastructure, which is supported by the National Science Foundation (Grant No. ECCS-2025462). Funding for CSMB is provided by the Office of Biological & Environmental Research in the Department of Energy’s Office of Science. A portion of this research used resources at the Spallation Neutron Source, a DOE Office of Science User Facility operated by the Oak Ridge National Laboratory. The authors gratefully acknowledge Prof. D.A. Tirrell and Prof. K. Zhang for Z_E_ and mCherry genes and AF-IQ *E. coli* and Dr. W. Leite for assistance with the SAXS experiments. We acknowledge the contributions of named and unnamed people whose health, lives, livelihoods, legacy, and privacy were extorted, often without compensation, consent, or regard to their safety, in the name of biomedical research. These men, women, and children were stripped of their humanity, and often their identity. We knowingly use resources and knowledge with the gratitude and respect not given previously. We commit to educating ourselves and others on the history and ethical failures of biomedical research, expressing our gratitude, and encouraging others to do the same.

Received: ((will be filled in by the editorial staff))

Revised: ((will be filled in by the editorial staff))

Published online: ((will be filled in by the editorial staff))

## Notes

### Competing Interest Statement

The authors have declared no competing interest.

