## Supplemental Information for "Intracellular biomacromolecule delivery by stimuli responsive protein vesicles loaded by hydrophobic ion pairing"

Table S1. ((Sequences of ELP variants. Subscripts in the ELP sequence column indicate number of repeats. Bolded amino acids are modifications to the original ELP sequence, in place of valines.))

| ELP Name | ELP Sequence |
| --- | --- |
| Y_5_-Z_R_-ELP | [VPGVG VPG**Y**G VPGFG VPGVG VPGVG]_5_ |
| H_15_-Z_R_-ELP | [VPG**H**G VPG**H**G VPGFG VPG**H**G VPGVG]_5_ |

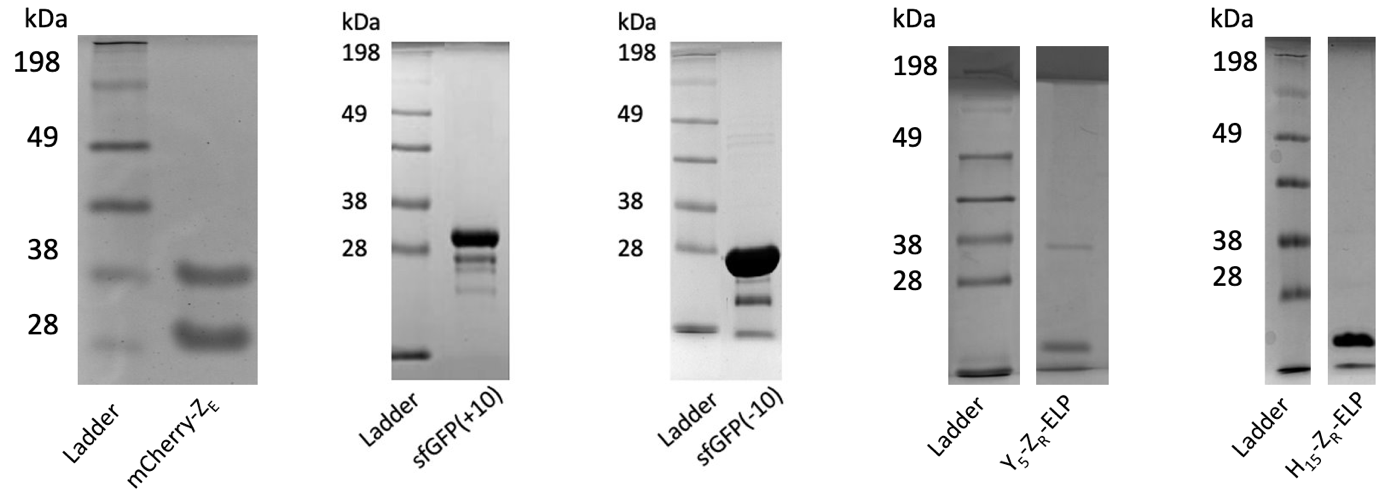

Figure S1. SDS-page gels of mCherry-Z_E_, sfGFP(-10), sfGFP(+10), Y_5_-Z_R_-ELP, and H_15_-Z_R_-ELP proteins. mCherry-ZE and the sfGFP samples when boiled results in multiple populations one for the folded fluorescent protein and the other for the partially folded protein.*^46^* The boiling hydrolyzes the N=C bond resulting in the decrease in fluorescence.*^49^* Y_5_-Z_R_-ELP has two bands where the lower band is protein monomer and the higher molecular weight band is the dimer. Dimers form due to the disulfide bonds between the terminal cysteine residues in each protein. The lowest band in Y_5_-Z_R_-ELP and H_15_-Z_R_-ELP gels is the dye front.

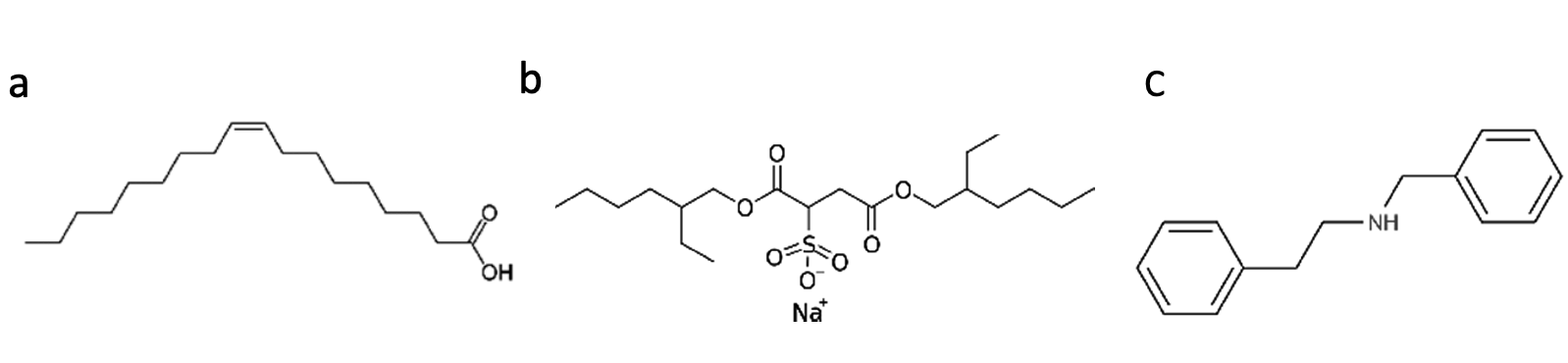

A

B

C

Figure S2. Structures of counterion (A) OA with a LogP=6.78, (B) SD with a LogP=5.2, and (C) BA with a LogP=3.6.^11^

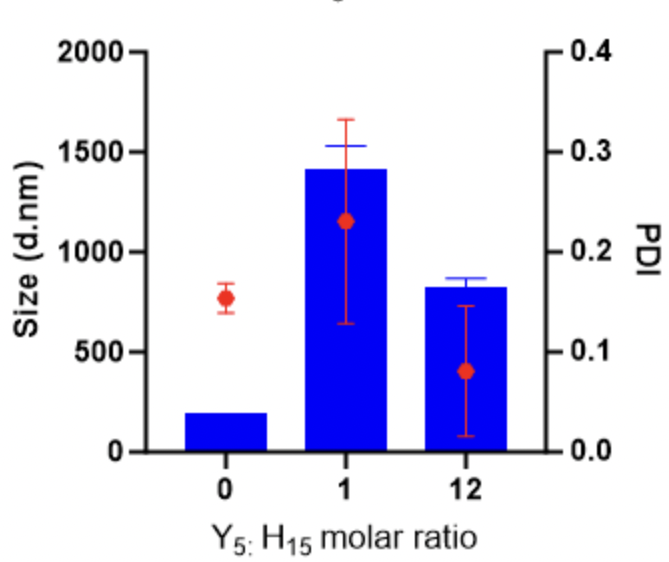

Figure S3. Protein vesicle hydrodynamic diameter and polydispersity index (PDI) of varying Y_5_-Z_R_-ELP: H_15_-Z_R_-ELP molar ratios. Blue bars represent size and red dots are PDI. Vesicles were composed of 0.15 M NaCl, 0.05 Z_E_/Z_R_ ratio, and 30 µM total ELP.

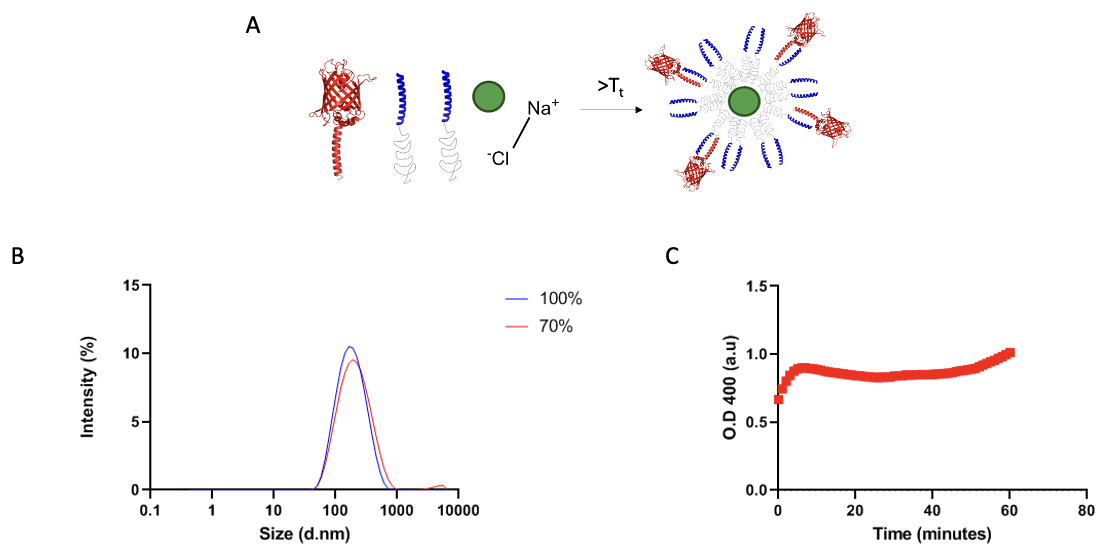
Figure S4. Turbidity profile of 0.15 M NaCl, 0.3 Z_E_/Z_R_ ratio, and 30 µM total ELP using a 12 Y_5_-Z_R_-ELP: 1 H_15_-Z_R_-ELP molar ratio solution over 1 hour after removing from ice to room temperature.

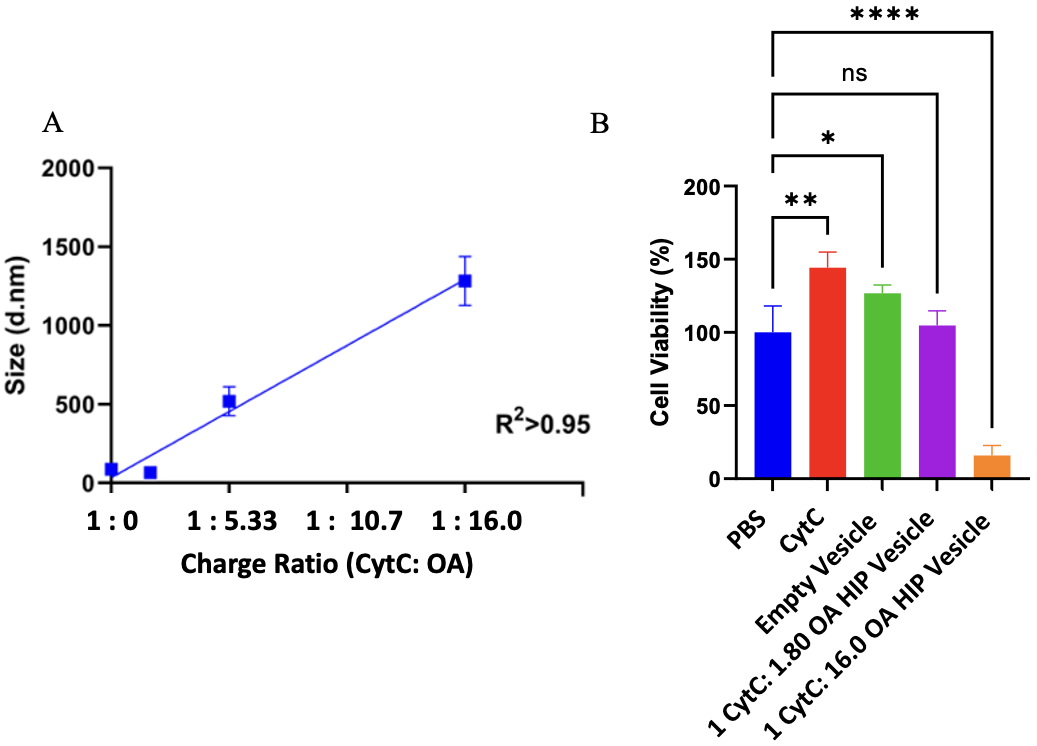
Figure S5. Characterization of HIP CytC OA vesicles with varying amounts of OA. (A) Hydrodynamic diameter of 0.15 M NaCl, 0.3 Z_E_/Z_R_ ratio, and 30 µM total ELP using a 12 Y_5_-Z_R_-ELP: 1 H_15_-Z_R_-ELP molar ratio solution protein vesicles loaded with the same concentration of CytC with varying concentrations of OA and (B) viability of 2D K562 cells treated with CytC vesicle groups including two mass ratios of HIP. Groups used 2 µM CytC. One-way ANOVA was utilized with p>0.05 n.s, p<0.05*, p<0.01**, p<0.0001****, and n=3 groups with each experiment repeated at least twice. Error bars are standard deviation from the mean.

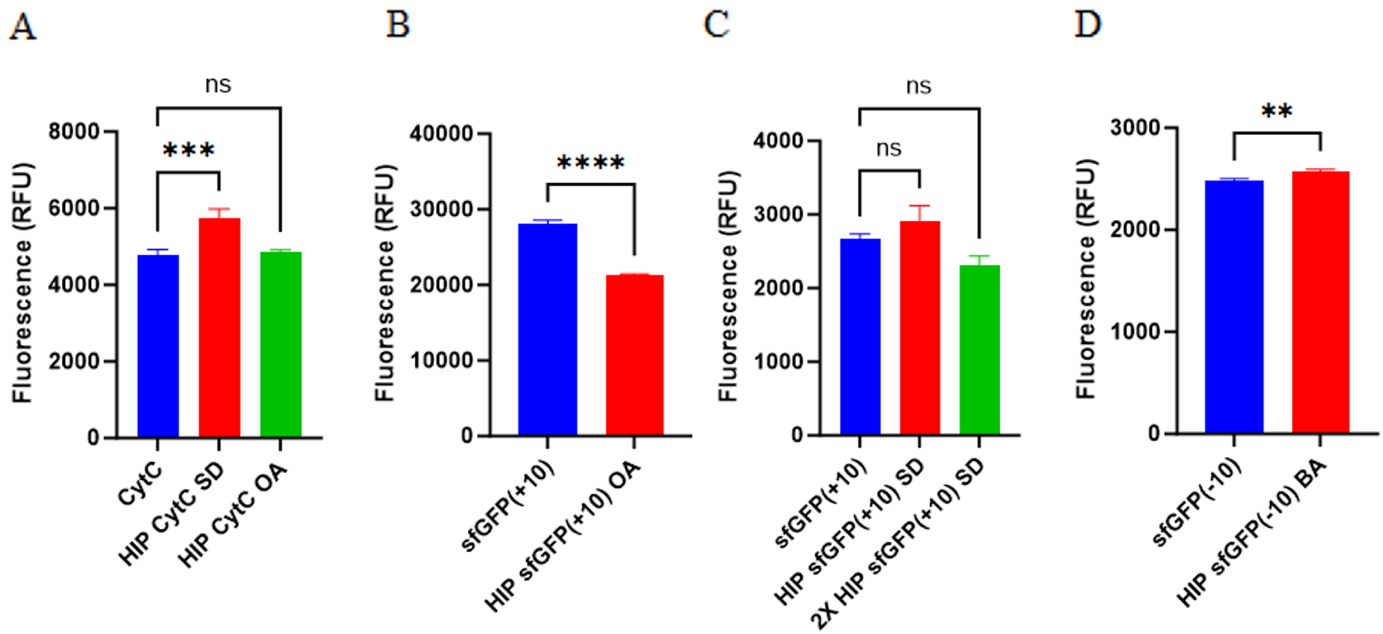

Figure S6. Counterion influence on (A) CytC, (B) sfGFP(+10) & (C) sfGFP(+10) and (D) sfGFP(-10) cargo fluorescence as a measure of change in protein structure. One-way ANOVA was utilized in (A) and (C) and a t-test was utilized in (B) and (D) with p>0.05 n.s, p<0.01**, p<0.001***, p<0.0001****, and n=3 groups with each experiment repeated at least twice. Error bars are standard deviation from the mean.

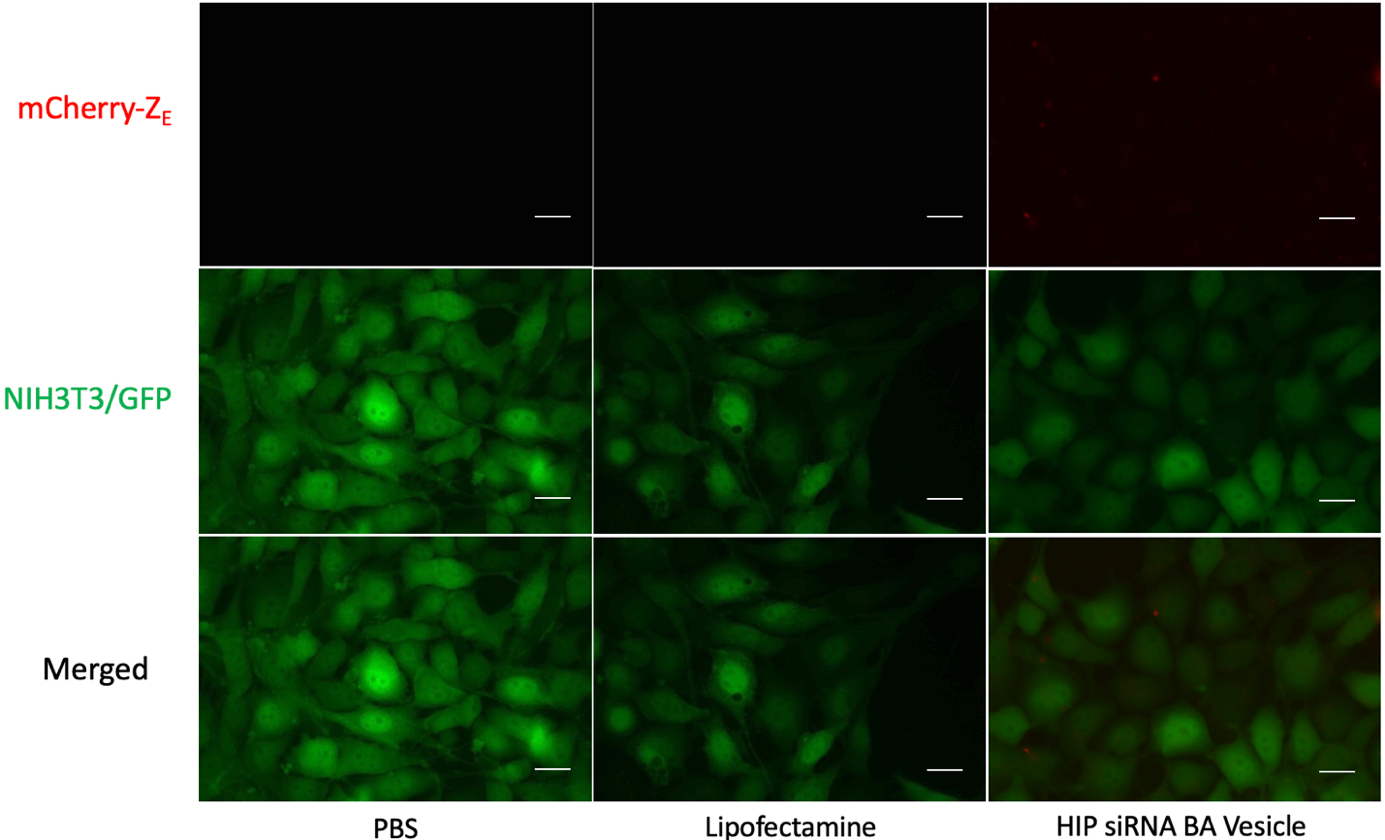

Figure S7. Epifluorescence images of GFP siRNA delivery to NIH3T3/GFP cells using pH-responsive protein vesicles after 48 hours. 150 nM siRNA was used for vesicle delivery while 100 nM siRNA was used for lipofectamine RNAiMAX. Vesicles were treated in the presence of serum-containing media while lipofectamine requires serum-free conditions. Scale bar is 5 microns.

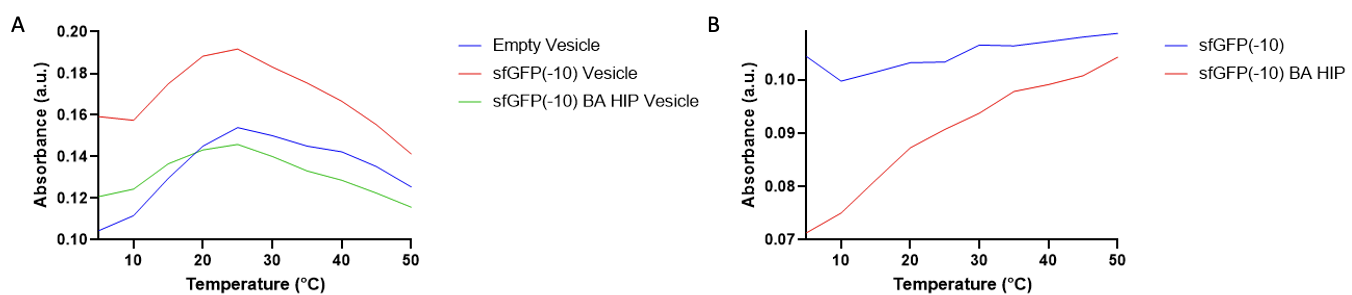

Figure S8. Transition temperature profiles of A) empty, sfGFP(-10), and sfGFP(-10) BA HIP vesicles with 10 µM sfGFP(-10) with a 1 degree/minute ramp rate. Transition temperature profiles of B) sfGFP(-10) and sfGFP(-10) BA HIP solutions with 10 µM sfGFP(-10) with a 1 degree/minute ramp rate.

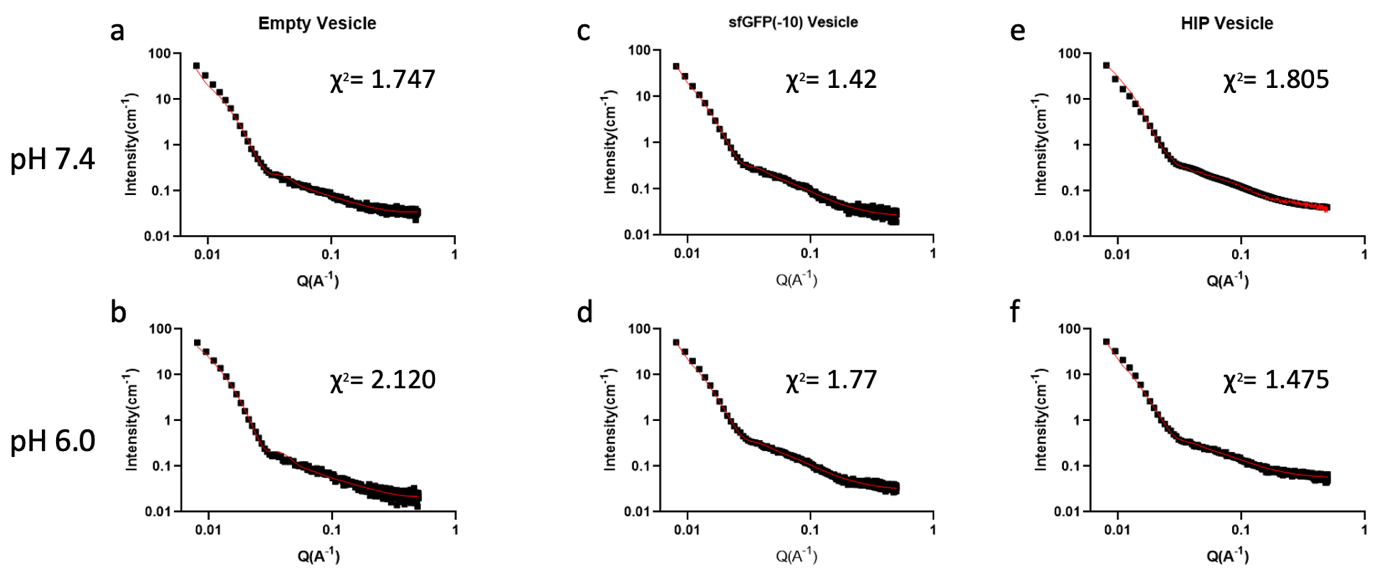

A

C

E

B

D

F

Figure S9. SAXS model fitting of empty protein vesicles (A) at pH 7.4 and (B) pH 6.0, (C) sfGFP(-10) vesicles at pH 7.4 and (D) pH 6.0, and (E) sfGFP(-10) BA HIP vesicles at pH 7.4 and (F) pH 6.0. Loaded groups each had 10 µM sfGFP(-10) and 0.816 mg/mL protein vesicle.

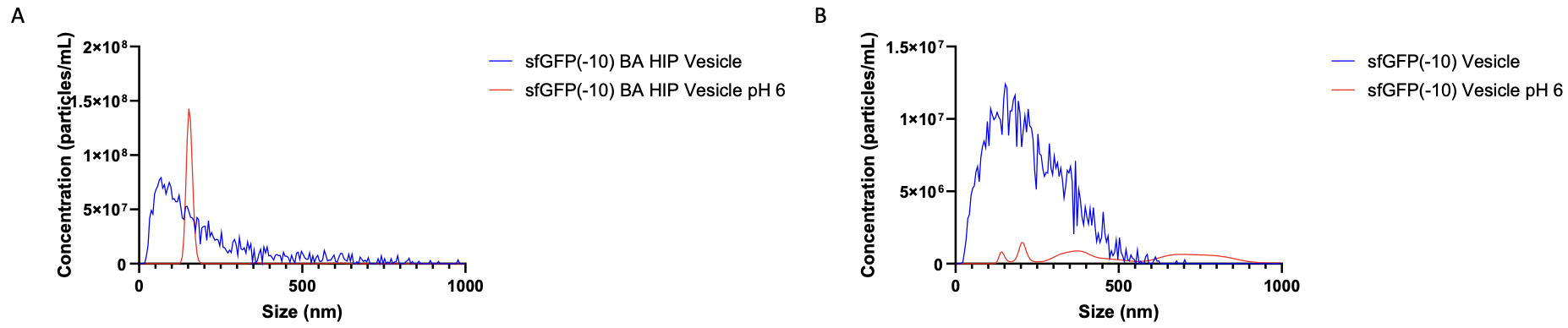

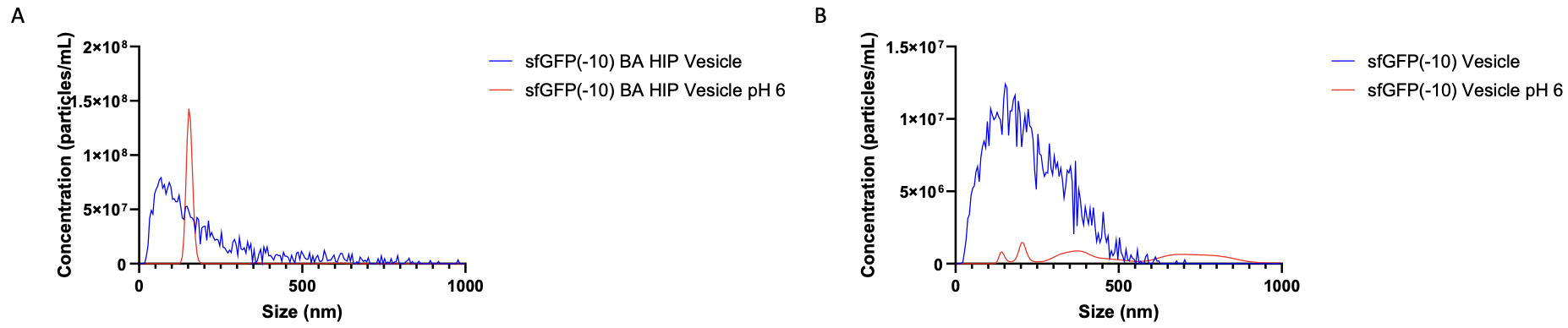

Figure S10. NTA analysis of loaded vesicles. (A) sfGFP(-10) and (B) sfGFP(-10) BA vesicles at pH 7.4 and pH 6. Vesicles were diluted by a factor of 10 to a total protein concentration of 0.0816 mg/mL.

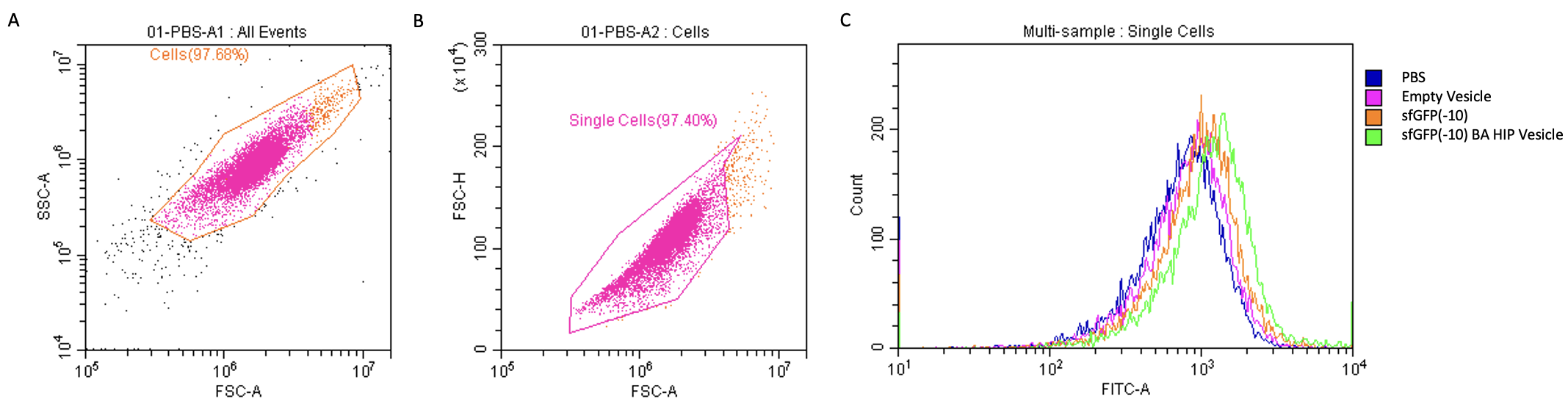

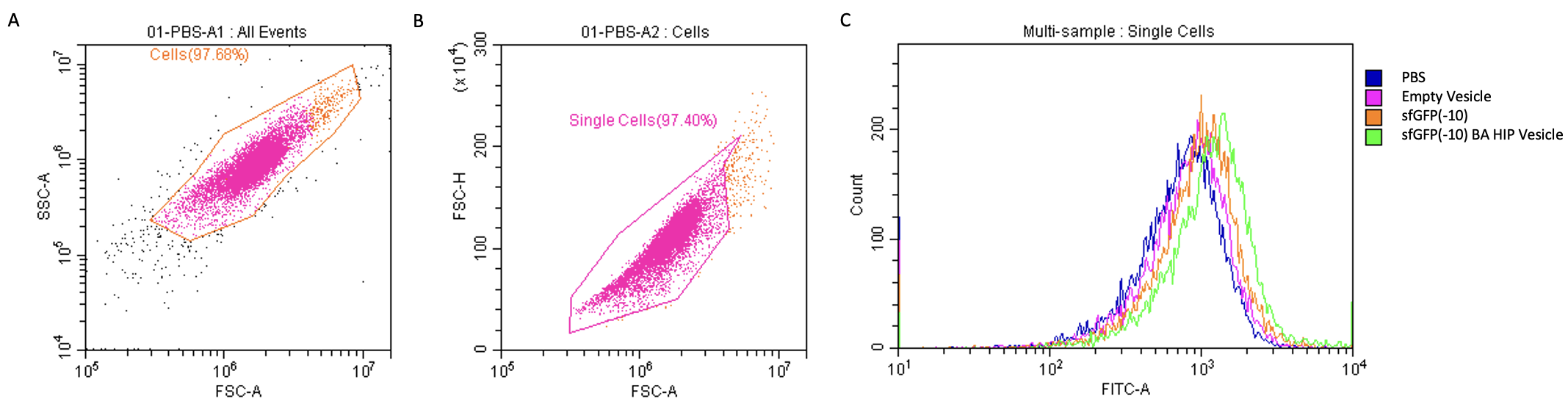

Figure S11. Example flow cytometry gating to select single cell populations. (A) Forward scatter area (FSC-A) and side scatter area (SSC-A) enable selection of cell populations. (B FSC-A and FSC height (FSC-H) enable selection of single cell populations by drawing tight gates to remove doublets. (C) Single cell populations were used to measure sfGFP uptake using median values.
